# Temperatures that sterilise males better predict global species distributions than lethal temperatures

**DOI:** 10.1101/2020.04.16.043265

**Authors:** Steven R Parratt, Benjamin S Walsh, Soeren Metelmann, Nicola White, Andri Manser, Amanda J Bretman, Ary A Hoffmann, Rhonda R Snook, Tom AR Price

## Abstract

Predicting how biodiversity will respond to increased temperatures caused by climate change is vital. However, our understanding of the traits that determine species’ response to thermal stress remains incomplete. Laboratory measurements of lethal temperatures have successfully been used to predict global species distributions and the vulnerability of species to future climate change. However, although it has long been known that fertility is sensitive to heat stress, temperatures that cause sterility have not been incorporated into predictions about how climate change will affect biodiversity. Here we show that male sterility temperatures predict the global distributions of 43 species of *Drosophila* substantially better than their lethal temperatures. This strongly suggests that thermal limits to reproduction can underpin how temperature affects species’ distributions. High temperatures impair male fertility across a broad range of animals and plants, so many organisms may be more vulnerable to high temperatures than currently expected.

## Main Text

We urgently need to understand how rises in temperature will impact biodiversity^1^. To do this the physiological, behavioral and evolutionary factors that underpin species’ current thermal distributions need to be considered^2,3^. Laboratory-derived estimates of the highest temperatures at which an organism can function provide measures of species’ thermal tolerances^5,6^. These measures of upper thermal limits have improved the accuracy of functional species distribution models^4^ which can be extrapolated to climate change scenarios to forecast future distributions^5^. Accurate predictions of species’ distributions are invaluable for prioritizing conservation efforts^6^ and predicting the invasion of pests and disease vectors^7^.

Upper thermal tolerance limits are usually based on the temperatures that cause loss of coordinated movement, coma, respiratory failure, or death: the species’ critical thermal limit. Despite these being measured in artificial laboratory conditions, critical limits correlate reasonably well with species’ distributions^8,9^ and have been used to estimate species’ capacity to tolerate temperature increases across their current range; their ‘thermal safety margins’^3,8^. However, critical limits can be higher than the temperatures that cause seasonal population declines in nature^10^. Therefore, population persistence may also be impacted by the effects of sub-lethal temperatures on traits such as reproduction. Sub-lethal temperatures cause losses in fertility in plants^11^, insects^12-14^, fish^15^, aquatic invertebrates^16^, birds^17^and mammals, including humans^18^. Thermal stress disrupts fertility by impacting physiological processes^12,17,19^ and influencing behavior and phenology^20^. The temperature at which fertility is lost, the thermal fertility limit (TFL)^20^, may be an important part of species’ true upper thermal limits. If TFLs are lower than critical limits, then many organisms will be more vulnerable to climate change than currently thought. If TFLs and critical limits correlate poorly then we may misidentify which species are most at risk from rising temperatures. We need to test whether TFLs measured in laboratory conditions can predict the distribution of natural populations better than measures of lethal limits.

## Comparing lethal and sterilizing temperatures

Fruit flies of the genus *Drosophila* have been a model taxon for thermal biology and biogeography^8,21^ due to their well understood biology, laboratory tractability, and the diversity of thermal environments the species inhabit. Here, we recorded three measures of upper thermal limits in adult males from 43 species of *Drosophila*. First, we compared fertility and survival limits under identical heat-stress conditions. We exposed flies to a 4-hour static heat stress at a range of temperatures from benign through to lethal. From these data we estimated both the temperature that is lethal to 80% of individuals (LT80), and the temperature at which 80% of surviving males are sterilized (TFL80). Measuring thermal traits under static temperature stress rather than slowly increasing temperatures (i.e. ramping) has received criticism^22^. However, ramping assays require an immediate observable response, such as flies losing coordinated motor function. Unfortunately, sterilization is not immediately observable, so we use static temperatures and assay fertility through subsequent matings. We score fertility at two time points: (i) cumulatively over 1-6 days post-heat, to capture any immediate sterilizing effect of heat, and (ii) 7-days after heat-stress to capture any recovery of fertility or delayed sterility. To compare our estimates of TFL80 and LT80 with a measure of lethal temperature under ramping thermal stress, we also assayed the CT_MAX_ of each species. This is the temperature at which males lose coordinated motor function under gradually increasing temperatures. CT_MAX_ is commonly used across animal systems to predict species’ sensitivity to thermal stress associated with climate change^3,8^.

We found that 11 of 43 species experience an 80% loss in fertility at cooler-than-lethal temperatures immediately following heat-stress (Extended Data Figure 1). Interestingly, rather than seeing a recovery of fertility over time, the impact of high temperatures on fertility was more pronounced 7-days post heat stress (Figure 1). Using this delayed measure of fertility, 44% of species (19/43) showed fertility loss at cooler-than-lethal temperatures. The difference between lethal and fertility limits ranged from 0°C to 4.3°C (mean of all species = 1.15 ± 0.22°C), and LT80 and TFL80 predict dramatically different relative ranking of species’ robustness to high temperature (Extended Data Figure 2). All three thermal limits significantly, positively correlate with each other (Extended Data Table 1). Despite deriving from different types of heat-stress, the correlation coefficient between CT_MAX_ and LT80 is larger than that between TFL80 and either measure of lethal temperature. Relatively low correlations between survival (measured as CT_MAX_ under dynamic conditions or LT80 under static stress) and fertility (measured under static heat stress) suggests that they are distinct phenomena, and that measuring both may be important for understanding how species respond to thermal stress.

**Figure 1:**
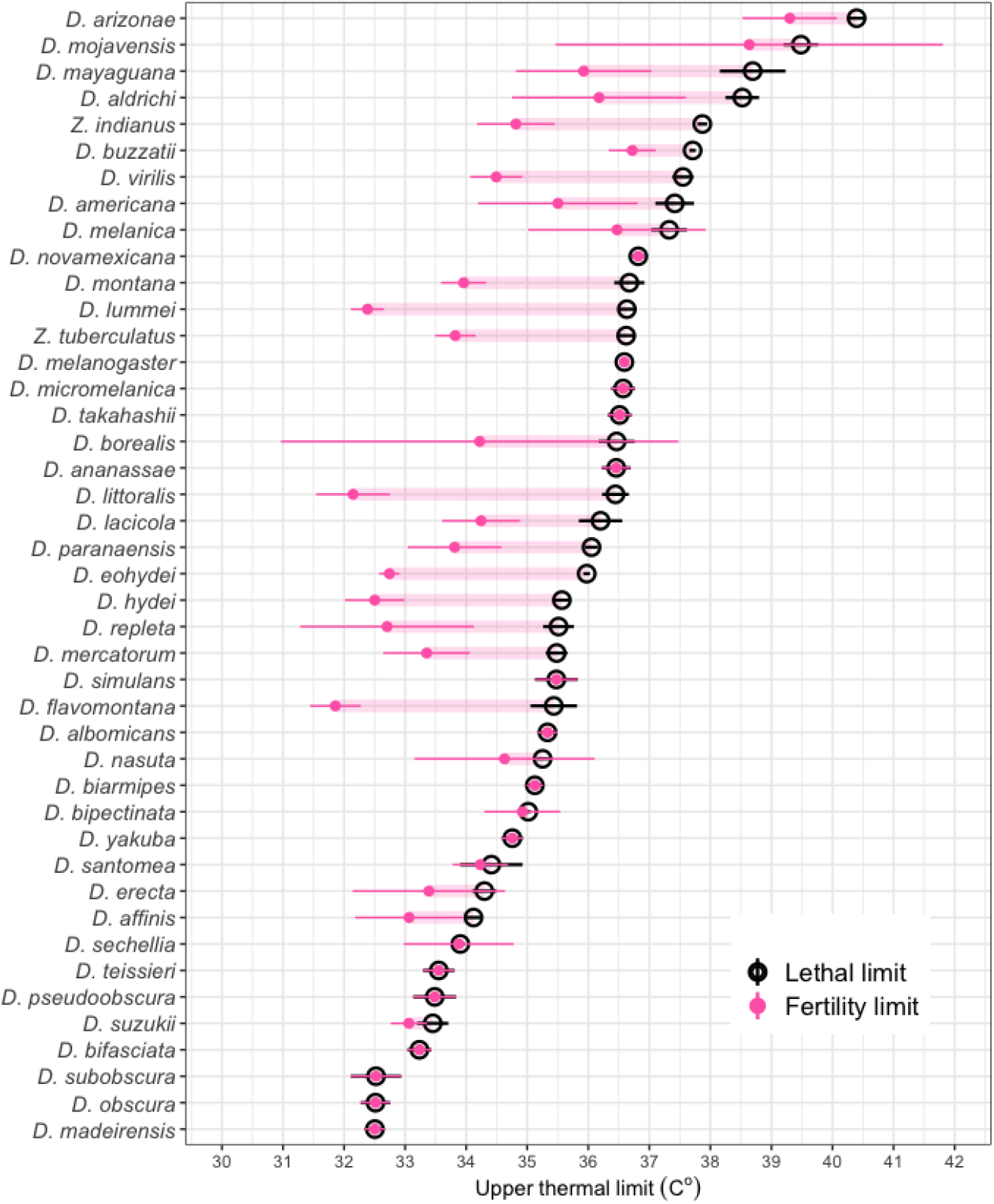
Upper lethal temperature (LT80, black circles) and upper thermal fertility limits (TFL80 measured 7-days after heat stress, pink points) of 43 species of *Drosophila*. 19 of 43 species show significantly lower thermal fertility limits than lethal limits. Species ranked by LT80 with estimated TFL80 joined by pink bar. We see considerable variation in TFL80 across the range of lethal values, with some species maintaining fertility at high temperatures. 95% CI are shown as error bars for both measures, differences between a species’ TFL80 and LT80 considered to be significant if these bars do not overlap. Our measure of TFL80 is capped at the LT80 to reflect ecological realism (see Methods). Fertility loss was measured seven days post heat stress (for fertility measured immediately following heat-stress see Extended Data Figure 1).

**Figure 2:**
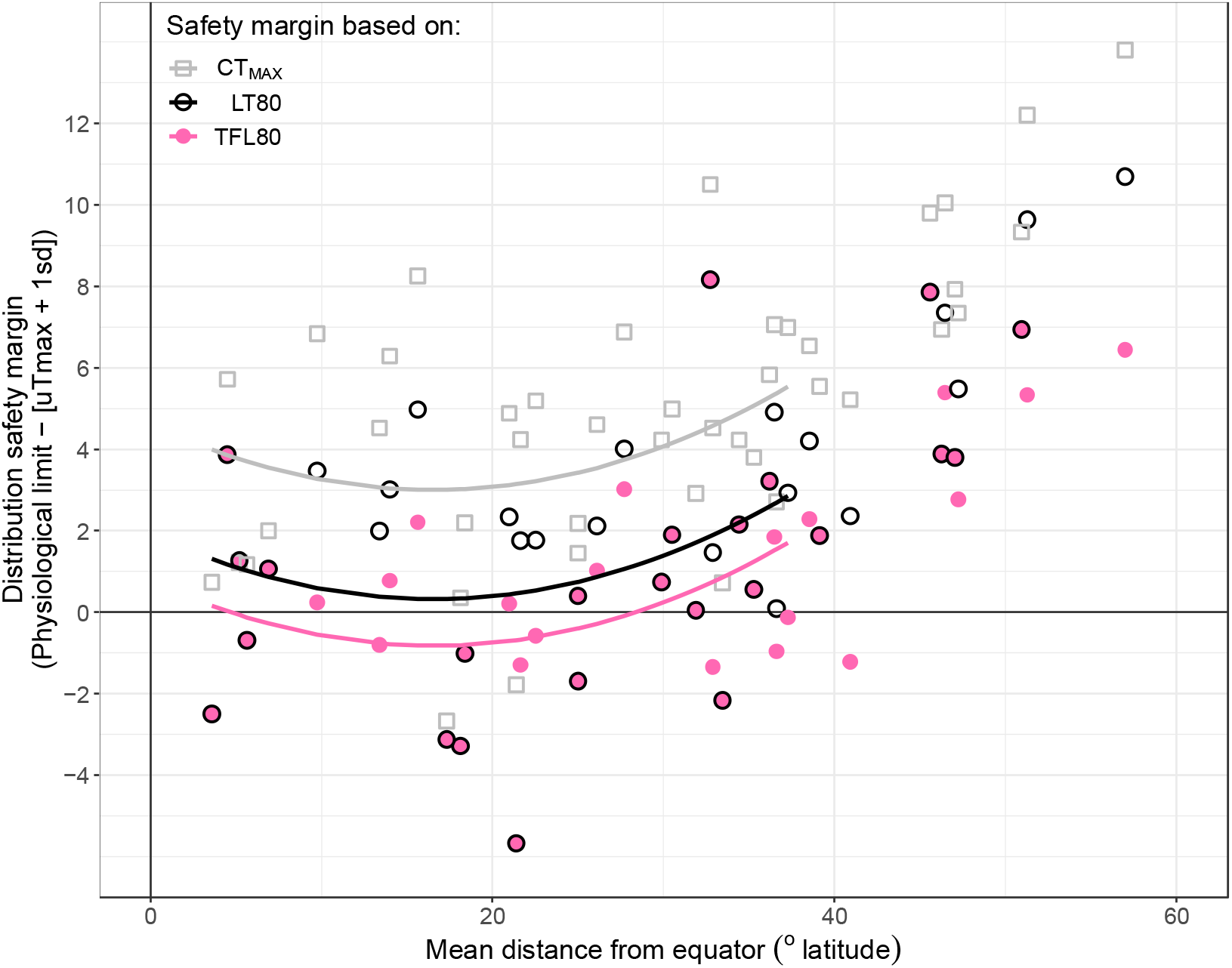
Distribution Thermal safety margins for 43 *Drosophila* species. Margins calculated using the physiological limit minus mean maximum air temperature + 1SD across all recorded locations for each species to capture populations at the boundary of the thermal range^8^. Grey squares and line are margins calculated with CT_MAX_, black circles and lines with LT80, and pink points and lines with TFL80 measured 7-days after heat stress. Points on or below the horizontal ‘0’ line represent species with populations that are currently exposed to sterilizing temperatures in the hottest part of the year. Fitted lines from models summarized in Extended Data Table 3. For central safety margins see extended Data Figure 3.

Our data confirm that fertility loss at sub-lethal temperatures is common across *Drosophila*. This suggests that studies based on lethal limits alone may overestimate the thermal tolerance of many species, which could be problematic for efforts to preserve rare and endangered species or control emerging pests and disease vectors. For example, we found that the crop pest species *Zaprionus indianus* (African fig fly) has a relatively high upper lethal limit (LT80 = 37.9°C) but a much cooler upper fertility limit (TFL80 = 34.8°C). This switches the species’ thermal hardiness rank from 5^th^ to 17^th^ of species here (Extended Data Figure 2). *Z. indianus* is currently expanding throughout the New World, and this disparity could have marked consequences for forecasting its putative range and hence where to concentrate control efforts^23^.

## Linking Thermal Limits to Distributions

Our work demonstrates that TFLs can be substantially lower than lethal temperatures, even when measured under the same conditions in the laboratory. However, the key question is whether TFLs are linked to organisms’ distributions in nature. To test this, we integrated existing distribution data of each sampled *Drosophila* species with global climate data. From this we estimated the mean maximum air temperatures species are likely to encounter where they occur in nature. Previous work using 95 *Drosophila* species found that the mean maximum summer air temperature measured in this way significantly correlated with CT_MAX_ of species from dry environments (<1000mm annual rainfall), but not for species from wetter environments^8^. We first recapitulated this finding in our smaller dataset of 43 species; our measurement of CT_MAX_ significantly predicts mean maximum environmental air temperature (PGLS: T_40_ = 2.647, *P* = 0.012) albeit this relationship negatively interacts with annual rainfall (PGLS: T_40_= −2.077, *P* = 0.044) and has reasonably low explanatory power (adjR^2^ = 0.186). We then tested how well our measurements of LT80 and TFL80 predict the mean maximum air temperature in species’ distribution ranges (Extended data Table 1). LT80 significantly predicted mean maximum environmental temperature (PGLS: T_40_ = 3.360, P = 0.002), but again explained less than 20% of the variation (adjR^2^ = 0.197). However, the relationship between TFL80 and mean environmental maximum temperature was stronger, both when TFL80 was measured immediately following heat-stress (PGLS: T_40_ = 4.225, P < 0.001, adjR^2^ =0.286) and 7 days later (PGLS: T_40_ = 5.014, P < 0.001, adjR^2^ = 0.365). We found no significant interaction with annual rainfall for either LT80 or TFL80. Comparing across all best–fit models, TFL80 measured 7-days after heat shock most strongly predicted mean maximum air temperatures in species’ environments, explaining 36.5% of the variation. This is an 85.3% and 95.8% improvement in accuracy (adjR^2^) compared to models with LT80 or CT_MAX_ respectively. Furthermore, we found qualitatively similar results when we used a more conservative 50% threshold for LT and TFL estimates (Extended Data Table 4). These analyses suggest that TFLs and species distributions are strongly linked in nature, and that fertility losses due to high temperature are an important determinant of where species occur.

Our approach based on TFL80 provides a significant improvement over current measures of upper thermal limits such as CT_MAX_, which have been valuable in predicting species’ responses to climate change^3,6,8,9^. Nevertheless, all laboratory measures of thermal limits have issues in reflecting ecological realism. They are optimized for high throughput of species and individuals, rather than to precisely replicate natural conditions, and often measure heat effects under artificial conditions. In our tests, we used some fly stocks that have been in culture for many generations and so may have phenotypes compromised by adaptation to benign laboratory temperatures or genetic drift. However, we find only weak evidence for lab adaptation in LT80 and no significant signal for lab adaptation in either CT_MAX_ or TFL80 (Methods and Supplementary Information). We also have not considered variability of physiological traits across different populations of a given species, and so cannot account for local adaptation. Despite these limitations, our measure of TFL does appear to be a useful measure, as it captures 37% of the variation in temperatures among species’ distributions.

## Predicting vulnerability to temperature

Predicting species’ vulnerability to current and future temperature extremes is critical to protect biodiversity^1^. “Thermal safety margins”, often measured as the difference between an organism’s physiological thermal limit and the maximum air temperature it is likely to experience may indicate the species’ vulnerability to climate change. Although safety margins can use complex measures of environmental temperature to improve accuracy (e.g. microhabitat availability, acclimation^24^, thermoregulation behavior of adults^3^, duration of exposure^24^), they remain reliant on comparable measures of thermal resistance across species and populations. We found that TFL80 measured seven days after heat-stress produced significantly smaller safety margins than either LT80 or CT_MAX_ (LMER: F = 28.23_2,84_, *P* < 0.001, Tukey multiple comparisons P <0.001, Extended Data Figure 3 & Table 3). For populations that already live near the upper edge of their thermal range, safety margins are reduced from a mean of 2.38 ± 3.46°C when based on LT80, to 1.23 ± 3.04°C when based on TFL80 (Figure 2). This predicts that 15 of the 43 species studied here have populations found in environments in which air temperatures exceed safety limits during the hottest part of the year. We illustrate the implications of this with case-study distribution models of *Drosophila flavomontana*; safety margins based on TFL80 predict a 17.9% reduction in habitable landscape compared to an identical LT80-based model under current climate conditions (Figure 3A). The disparity between predictions based on sterility and lethality grew to 48.0% by the year 2080 under moderately optimistic future climate forecasts (ICCP-AR5 RCP 4.5, Figure 3B), and to 58.9% under pessimistic climate change scenarios (ICCP-AR5 RCP 8.5, Figure 3C). TFL-based models also predict that by 2080 the available habitat for *D. flavomontana* will have reduced by 42.3% and 62.9% under RCP4.5 and RCP8.5 respectively.

**Figure 3:**
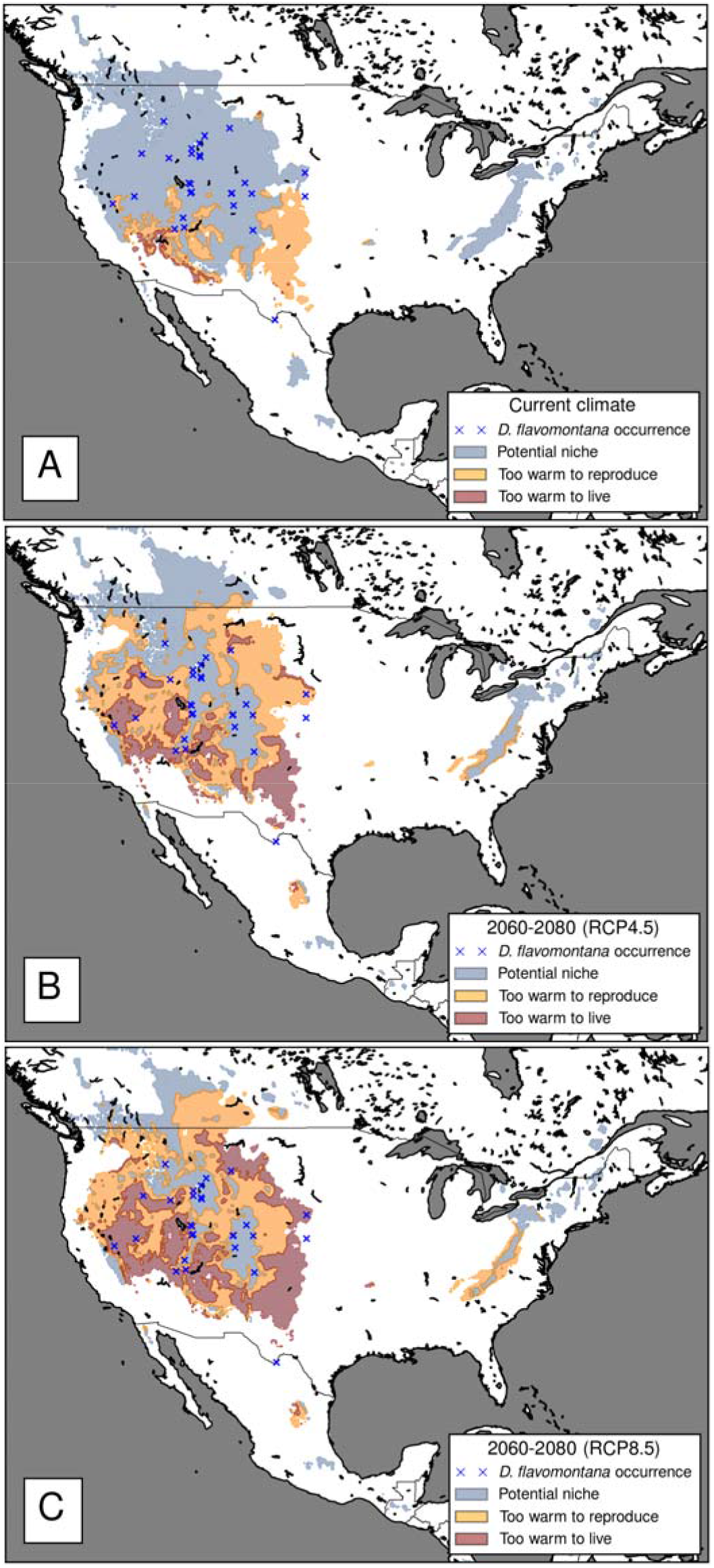
Potential habitat range of *Drosophila* flavomontana (LT80 = 35.4°C, TFL80 = 31.9°C) under A) current and B & C) possible future climate scenarios (B = RCP4.5 ‘moderately optimistic’, C = RCP8.5 ‘pessimistic’, predicted for 2060 - 2080). Colored areas in each panel represent suitable habitat range predicted by the model that excludes maximum temperature. Red areas show regions where maximum summer temperatures exceed LT80. Orange areas show regions where maximum summertime temperatures exceed TFL80. Blue regions are areas where limits for *D. flavomontana* are not exceeded all year.

If our data for *Drosophila* can be extrapolated to other organisms, then male fertility losses at high temperatures may be common, occurring at substantially lower temperatures than the species’ upper lethal limit. The limited data on fertility at extreme temperatures supports this, with losses in male fertility at high temperatures seen across a broad diversity of organisms^20^. Even species that might be predicted to have high thermal tolerance can show evidence of thermal fertility losses. For instance, the zebra finch, a high temperature adapted desert-dwelling endothermic species with naturally high body temperature and good thermoregulation, shows substantial damage to sperm at temperatures it regularly experiences in nature^17^. Behavioral thermoregulation could potentially reduce the impact of high temperatures on fertility in nature. However, while some evidence suggests that *Drosophila* are able to behaviorally thermoregulate in the lab^25^, evidence for flies choosing cooler microclimates such as leaf litter, shade, or higher altitudes in nature is mixed^26^. Further, many species are able to survive high temperature periods by aestivating as adults, eggs or pupae. This may explain why our data predict negative thermal safety margins for some species. Despite these potential mechanisms, we still find that species’ distributions are predicted by thermal fertility limits.

Our work emphasizes that temperature-driven fertility losses may be a major threat to biodiversity during climate change. We urgently need to understand the range of organisms likely to suffer similar fertility losses in nature, and the traits that predict vulnerability. However, we currently do not understand the physiology underlying variation in TFLs between species, nor the selective forces that created this variation. Ultimately, we need to know whether evolution for higher TFLs will allow species to adapt to a warming environment.

## Supporting information

Supplementary Methods, Results and Tabls of Species details

## Materials and Methods

### Animal maintenance

*Drosophila* species were obtained from stock centers, other research groups, or field collections (Supplementary Information Table 1). All species were kept in temperature-controlled rooms at temperatures selected to avoid effects of heat-hardening or stress (Supplementary Information Table 1). All rearing temperatures were well below those observed to be stressful, indeed our warmest rearing temperature (25°C) was several degrees cooler than the coolest lethal temperature we observe for any species (32°C). All stocks were kept at moderate density (20 – 50 breeding pairs per vial or 50 – 100 pairs per bottle culture) at 12:12 L:D and ambient humidity. Cultures were tipped to fresh food every 7 days and a new generation was made every 4 – 6 weeks depending on the speed of species development. Species were reared on one of four food types selected to minimize heterogeneous nutritional stress; A = ASG (10g agar, 85g Sucrose, 20g yeast extract, 60g maize, 1000ml H_2_O, 25ml 10% Nipagin), B = Banana (10g agar, 30g Yeast extract, 150g pulped banana, 50g Molasses, 30g Malt Extract, 25ml 10% Nipagin, 1000ml H_2_O), P = Propionic (10g Agar, 20g Yeast extract, 70g cornmeal, 10g soya flour, 80g Malt Extract, 22g Molasses, 14ml 10% nipagin, 6ml Propionic acid, 1000ml H_2_O), M = Malt (10g agar, 20g Yeast, 60g Semolina, 80g Malt, 14ml 10% Nipagin, 5ml Propionic acid, 1000ml H_2_O).

### Measuring upper thermal limits

We assayed three metrics of upper thermal limits in sexually mature males from 43 species of *Drosophila*: Lethal Temperature (LT), Thermal Fertility Limit (TFL) and Maximum Critical Temperature (CT_MAX_). We tested these limits in only males because of the substantial evidence that male fertility is particularly vulnerable to high temperatures^20^. All experiments were blinded by numbering vials randomly across treatments.

### Measuring Lethal Temperature (LT) & Thermal Fertility Limit (TFL)

We measured LT and TFL under static temperature conditions by allocating groups of flies to separate four-hour temperature pulses and recording their survival and fertility. Using static temperatures to measure thermal threshold traits has received criticism^22^. However, the use of ramping temperatures requires an immediate observable response, such as flies losing their coordinated motor function, which allows the researcher to visually determine the temperature at which the function was lost. Unfortunately, in *Drosophila* fertility is internal and has no visible marker, so there is no directly observable response that would indicate that a male has become sterile as temperatures increase. Hence, we have to use static temperatures and assay fertility by subsequently mating the male to females. Measuring LT under static temperature stress allows us to directly compare measures of fertility loss and lethality under identical heating conditions.

Newly eclosed virgin adult males were collected into plastic vials in groups of 10 over a 48-hour period and allowed to sexually mature for 7 – 21 days depending on species (see Supplementary Information Table 1 for maturation times). Mature males were transferred to fresh vials with standard ASG food and a cotton-wool stopper and allocated to 3D-printed floating racks in pre-heated waterbaths (Grant TXF200) set to a range of temperatures (Supplementary Information Table 1). ASG food was used during heating for all species because preliminary work had shown that the survival under heat stress was influenced by food type. We monitored the temperature during the assay in a fresh ASG vial without flies with a k-type thermocouple attached to a data logger (Pico Technology TC-08). Males were heated for 4 hours between ∼10am - ∼2pm and then returned to temperature-controlled rooms set to the species’ benign temperature. We scored survival of males the next morning to account for any immediate recovery or delayed death. We scored flies as ‘dead’ if they were completely immobile or had become stuck on their backs in the fly food – the latter would indicate that the flies were completely incapacitated which would likely be lethal in the wild. Surviving males were individually aspirated into separate vials containing their designated food type and 3-4 sexually mature virgin females. Males remained in these vials at their benign temperature and were allowed to mate freely for 6 days. After 6 days males were transferred to a second vial with 1-2 more virgin females and allowed to mate for 24 hours. Giving males two sets of females to mate with allows us to score fertility at two time points to capture any recovery or delayed sterilizing effect of heat over time. Females in both sets of vials were left to lay eggs for 3-5 days (depending on fecundity of the species) after the male had been removed. Vials were scored for the presence of larvae or their distinctive track marks in the food. We found no significant differences in mortality between heat-stressed and control males during the 1-week oviposition period, so we do not measure LT multiple time points.

We used the ‘drm()‘ function in the ‘drc’ package in R to fit dose response curves to survival and fertility data across temperatures for each species. The models generate point estimates of the temperature required to reduce survival or fertility by a given percentage relative to the maximum value in the data. We estimate thresholds based on 80% reduction (LT80 and TFL80 hereafter) because this is likely to represent a substantial threat to population stability for most species. To test the robustness of our findings we repeated all analyses with more conservative 50% reduction estimates (LT50 and TFL50 hereafter) and present these results in the Extended Data (Extended Data Table 3). We used 3-parameter versions of the log-logistic model which fixes the lower limit of the predicted curve to 0 to reflect that we expect fertility and survival to eventually reach 0 over an infinite range of temperatures. In these models we do not consider dead males to be sterile to avoid statistically confounding the effects of heat stress on survival and fertility. We only allow TFL80 to be lower than or equal to the species’ LT80 because estimating 80% fertility loss in populations that have already experienced >80% mortality provides little informative data. Furthermore, we only consider a species’ TFL to be statistically lower than its LT if the 95% confidence intervals of these two point-estimates do not overlap.

In total, 14742 males were exposed to heat treatment, of which 10925 survived and 9064 went on to be tested for heat-induced sterility. The mean sample size at each temperature per species was 36.5 ± 0.5sem, ranging from 10 – 88 individuals. Because fertility is inherently only measurable in males that survive heat stress, we may be measuring TFLs in more thermally hardy individuals – i.e. those that survive. In the context of our study, this will make our estimate of any difference between LTs and TFLs more conservative.

### Measuring CT_MAX_

Studies typically estimate upper thermal limits under dynamically ramping heat conditions^6,8,24,27^, rather than under static heat-stress as we have used here for TFL and LT. To compare our static measures of TFL and LT with patterns of heat tolerance found using the more widely used dynamic assays, we measured upper critical limits of our 43 *Drosophila* species under gradually ramping heat conditions – their critical thermal maxima (CT_MAX_). This allows us to compare how static and dynamic measures of critical heat tolerance predict species’ distributions.

We measure CT_MAX_ as the mean temperature at which male flies lose coordinated movement and are unable to right themselves. Sexually mature males were anesthetized with CO_2_ and isolated into individual glass sample tubes sealed with a rubber stopper. Sample tubes were left at ambient room temperature for at least 30 minutes to allow males to fully recover from CO_2_ anesthetization. Preliminary trials found no difference in CT_MAX_ between anesthetized and non-anesthetized male *D. melanogaster*. Sample tubes were then submerged in a water-filled glass aquarium held at the rearing temperature for the species (18, 23 or 25°C) by a heated circulator (Grant TXF200). The temperature was then increased at a rate of 0.1°C per minute and the temperature at which flies collapsed for 30 seconds and did not right themselves after being disturbed (tapping the vial) was recorded. The temperature of the waterbath was continuously recorded independently of the heating unit with a data logger (Elitech RC-61). The mean sample size for each species for CT_MAX_ assays was 19.53 ± 1.77.

Genetic and physiological changes due to lab domestication have been reported in a number of insects^28-30^. Because the 43 species of *Drosophila* we use here varied in the time spent in lab culture we tested for confounding effects of lab adaptation on physiological thermal limits (Supplementary Material). We found only minor signals of lab adaptation in measures of LT80, and no significant signal in either CT_MAX_ or TFL80 (Supplementary Information). We see no strong evidence that lab adaptation has degraded inter-species variation in thermal traits. Further, we can replicate our core results with a subset of the 11 most recently caught species (Supplementary Material). This supports work in other dipteran systems that have shown long-term lab stocks to be good proxies for thermal traits in natural populations^31^.

### Repeatability

It was impossible to assay upper limits across all 43 species simultaneously due to the number of flies that could be processed simultaneously, and the different temperature ranges required for each species. Further, inter-species variation in development time prohibited us from synchronizing all 43 species to run the assay in a completely randomized block design. Instead, we independently assayed CT_MAX_, TFL and LT for each species between June 2018 – November 2019.

To test the repeatability of our measure of LT80 we ran a single control block in February 2020 with 22 species that produced enough sexually mature virgin males in synchrony. We simultaneously exposed these flies to a range of 6 temperatures that covered all species’ estimated LT80. We then tested the strength of the correlation between our original LT80 estimates and those given by this replicate block. We determined if any given species showed significantly different estimates based on whether the confidence intervals of the two runs overlapped (Extended Data Figure 4). Sample sizes for this control run ranged from 3 – 10 males per temperature per species, and involved 1152 male flies in total.

To test repeatability for our TFL assay we ran two independent blocks of the full assay on *Drosophila virilis* and compared the TFL80 estimate. The blocks were run 6 months apart on the same equipment and fly stock, and data were collected by two separate researchers. We compare the estimated TFL80 from each run and consider differences to be significant if the 95% confidence intervals do not overlap (Extended Data Figure 5).

Repeatability of CT_MAX_ was estimated by randomly allocating species from the same rearing temperatures into blocks of 3 – 5 species and measuring their mean CT_MAX_ as described above. We compared mean CT_MAX_ values with those obtained when species were run individually (Extended Data Figure 6).

### Correlations between upper thermal limits

We explored the correlation between all three physiological limits to test if TFL captures additional differences in species’ thermal hardiness. We used several parallel approaches to test the strength of correlations. We accounted for evolutionary relationships between species by including information from a *Drosophila* phylogeny published by Patrik Röhner and colleagues^32^. Firstly, we calculated phylogenetically independent contrasts (PICs) for each trait and correlated these using standard linear models forced through the origin at 0. This approach assumes that each trait completely evolves along a Brownian model of evolution. Second, we analyzed each pair of traits with phylogenetically controlled least squares regression (‘pgls’ in the R package ‘caper‘) in which we estimate the phylogenetic signal (Pagels λ) in model residuals with a maximum likelihood approach. This approach is sensitive to the order of x and y variables, so we ran both inversions of each pairwise correlation. Finally, we used the ‘corphylo’ function in the ‘ape’ package to generate a correlation matrix of all three traits in which phylogenetic signal for each is estimated assuming an Ornstein-Uhlenbeck process. We also compared the output of these phylogenetically controlled approaches to standard linear models. In all analyses we used TFL80 measured 7-days post heat stress because we found that the sterilizing effect of temperature is delayed in many species.

### Species distributions and environmental variables

Geographic distribution data for each species were obtained manually from TaxoDros.uzh.ch as coordinates and location names. Location data was systematically cleaned with functions in the ‘CoordinateCleaner’ R package to check for appropriate decimalization, remove invalid coordinates, remove country centroids, remove locations over water, remove locations centered on biodiversity institutions, remove location with equal longitude and latitude, and flag outliers. Remaining coordinates were then plotted on maps and outliers were investigated in the primary literature and removed or corrected when appropriate. To minimize spatially biased oversampling we then used the ‘thin’ function in ‘spThin’ R package to reduce the dataset down so that no two points for a given species occur within 10km of each other. We ran this independently for every species except three small island endemics (*D. santomea, D. sechellia* and *D. madierensis*) to preserve variation in their sparse distribution records.

The cleaned dataset of geolocations was integrated with rasterized bioclimate data from the WorldClim V2.0 database using functions from the ‘Raster’ package in R^33^. Our intention was not to fully reassess multiple climate variables as predictors for *Drosophila* distributions, because this has received extensive attention in the literature^8,24^. Instead, we tested if TFLs correlate and interact with variables previously identified as important for predicting physiological limits and distribution in *Drosophila*; the maximum temperature reached in the hottest summer month (WorldClim Bio5 – T_max_ hereafter) and annual precipitation (WorldClim Bio12 - P_ANN_ hereafter). Mean and standard deviation of each climate variable for each species were calculated across all global locations.

### Comparing fit of CT_MAX_, LT and TFL to species’ thermal habitats

We used phylogenetically controlled linear models to test the relationship between each physiological limit and maximum air temperatures found in species’ natural environment (T_max_). We analysed our data with T_max_ as the response variable because previous work has found that species’ phylogenetically constrained physiology determines where they occur more readily than local adaptation drives phenotypic change^8^. Because of the strong correlations between the three measures of upper thermal limits we could not include them all as predictors in the same model with T_max_ as the dependent variable. Instead, we fit separate models with each of TFL80, LT80 or CT_MAX_ as independent predictors of species’ T_max_ and compare the goodness of fit. We included annual precipitation (P_ANN_) as an interacting covariate with each thermal limit because previous work has found that the relationship between lethal limits and T_max_ degrades for species that occur in more humid habitats^8^. We used AICc scores and visualization of residuals to retain or drop the interaction term from models. We do not retain P_ANN_ as a main term because we are not testing if spatial patterns of temperature correlate with spatial patterns of rainfall independently of *Drosophila* biology. All predictors are centered and scaled to the mean. We compared the adjusted R^2^ of each best-fit model for each upper limit.

### Calculating and modelling thermal safety margins

Thermal safety margins represent the physiological capacity of organisms to tolerate temperature increases in their native distribution. As per Kellermann (2012)^8^, we calculated this in two ways: “Central Safety Margins” are the physiological limit minus the mean T_max_ for each species, “Distribution Safety Margins” are the physiological limit minus the mean T_max_ + 1 standard deviation. “Distribution” safety margins have been argued to account for populations of species that live at the upper thermal range of species’ distribution. Safety margins were calculated separately using either CT_MAX_, TFL80 or LT80 as the physiological limit.

We tested for significant differences between safety margins calculated with each physiological measure with linear mixed models using the ‘lme4’ package in R. We set ‘physiological limit’ (CT_MAX_, TFL80 or LT80) and mean absolute latitude of each species known distribution locations as interacting predictors. Latitude was fit as a curvilinear (latitude^2^) predictor following visualization of the raw data and because previous studies that have found curvilinear patterns in safety margins over latitudinal range^3,8,34^. We included species identity as a random intercept to account for non-independence between safety margin values calculated for TFL80, LT80 and CT_MAX_ for each species. We used AICc and visualization of model residuals to reach a minimum best-fit model through stepwise selection. We used Type II Wald’s F test in the ‘car’ package to determine the significance of main effects in final models. We used post-hoc Tukey contrasts implemented with the ‘emmeans’ package to detect significant differences between safety margins based on the three physiological limits. “Central” and “Distribution” safety margins were modelled separately.

### Predicting current and putative future distributions of *D. flavomontana*

As a case-study demonstration of the importance of TFLs, we used MaxEnt v3.4.1^35^ to model potential distributions of *D. flavomontana*. Occurrence data from TaxoDros.uzh.ch were further slimmed so that points within 25km of each other were clustered together (N = 24). First, we chose climatic and other variables to predict a potential distribution of *D. flavomontana* that is independent of the upper thermal limit. After excluding variables that strongly correlated or had little to no impact on the model results (<3% contribution), we used four of the tested 15 variables: mean winter temperature (WC Bio 11, from the WorldClim2.0 database), precipitation seasonality (WC Bio 15), an agricultural index^36^, and elevation^37^. We then ran 10 cross-validations with maximum iterations of 5000, using 10000 random pseudo-absences. To distinguish suitable from unsuitable regions, we used the threshold that maximized Youden’s index (maximum training sensitivity plus specificity)^38^.

We then constrained the suitable area by the upper physiological thermal limits measured above: i) the lethal temperature measured as LT80 (35.4°C), and ii) the sterilizing temperature measured as absolute TFL80 (31.9°C). We use the maximum temperature of the warmest month as reference (T_max_). Using T_max_ directly as a predictor variable for *D. flavomontana* had close to no effect on the distribution (0.1% contribution).

For future predictions, we chose two carbon emission scenarios from the IPCC-AR5: RCP4.5 (moderately optimistic) and RCP8.5 (pessimistic). We used precipitation and temperature predictions of the NorESM1-M model from the NEX-NASA GDDP data set^39^, as this model produces a median trend in terms of temperature increase^40^. The agricultural index and elevation had to be used as present day data. Predicting and constraining the possible future distribution followed the same procedure as for the current distribution.

### Data and Code Availability

All data and analysis code required to recreate this study will be made freely available on Dryad upon acceptance of the paper.

## Acknowledgements

Natasha Mannion and Dr Angela Sims for assistance with experiments, Dr Ben Longdon and Dr Katherine Roberts for flies, Rowan Connell for designing 3D-printed equipment. Funding: NERC grant NE/P002692/1 to TP, AB, AH & RS. SNF P300PA_177830 to AM. NIHR HPRU EZI to SM.

## Author Contributions

Conception: TP, AB, RS, AH, SP. Methodology & data collection: SP, BW, NW. Data curation: SP & SM. Analysis: SP, AM & SM. Original Draft: SP, TP, AB, RS & AH. Review & Editing: All authors.

## Data and materials availability

All data and analysis R code will be deposited on Dryad upon acceptance of this manuscript.

## Competing Interest

The authors have no competing interest to declare.

**Supplementary Information** is available for this paper, including Supplementary Table 1 – Details of species used.

## Extended Data Legends

**Extended Data Figure 1: TFL80 and LT80 of *Drosophila* immediately after heat stress**. Several species of *Drosophila* lose 80% fertility at cooler-than-lethal temperatures immediately following heat-shock. LT80 (black circles) and TFL80 (pink circles). Errors for both measures are 95% confidence intervals generated from dose response model estimates. Fertility loss measured as ability to sire any offspring between 1 – 6 days post heat-stress.

**Extended Data Figure 2: Relative ranking of species thermal robustness based on LT80 and TFL80**. Most to least heat tolerant species ranked from top to bottom. Pink bars indicate species with significantly lower TFL80 than LT80. Data based on TFL80 recorded 7-days post-heat shock.

**Extended Data Figure 3: Central Thermal safety margins for 43 *Drosophila* species**. Margins calculated using the physiological limit minus the mean maximum temperature across all recorded locations for each species. Grey squares and line are safety margins calculated using CT_MAX_ values, black circles and lines are based on LT80, and pink points and lines are based on TFL80 measured 7-days after heat stress. The three physiological limits produce significantly different safety margin estimates (Central safety margins: lmer: F_2_ = 28.23, P < 0.001, estimate = −1.17). TFL80 measured at 7 days post-heat stress predicts mean central safety margins of 4.97 ± 2.42°C. This is significantly smaller than margins based on CT_MAX_ (mean central safety margins: 8.81 ± 3.26°C, pairwise Tukey tests P<0.001) and LT80 (central safety margins: 6.13 ± 3.09°C, pairwise Tukey tests P<0.001). Latitude shown here is the mean absolute latitude i.e. the sign of negative latitudes has been removed to give relative distance from the equator. Fitted lines are predictions from models in which the limit type (TFL, LT or CT_MAX_) and latitude^2^ are fixed effects and species identity is a random effect to account for repeated measures across species (see Extended Data Table 3).

**Extended Data Figure 4: Repeatability of LT80**. Repeatability was high across the 22 *Drosophila* species tested. Left panel: There was no significant difference in the estimate of LT80 between full runs and control repeats in any of these species (significance measured as non-overlapping CIs of point estimates of LT80). Pink points = estimate from single species assay, gold point = estimate from simultaneous multispecies control assay. Right panel: The correlation between the two independent measurements of LT80 across these species was strongly positive and explained a high degree of variation in the data (coefficient = 1.06, R^2^ = 0.96).

**Extended Data Figure 5: Repeatability of TFL80 for *Drosophila virilis***. We independently repeated the TFL80 assay for one species in which we found a significant difference between LT80 and TFL80. Two independent runs of sexually mature males were conducted by two researchers 6 months apart. Red = original data used to calculate TFL80 for *D. virilis* in this manuscript, blue = repeated assay. Dashed lines intersect at 80% threshold. Line fits predicted by 3-parameter dose-response model. The fly stock and equipment were identical.

**Extended Data Figure 6: Repeatability of knockdown CT**_**MAX**_. Left panel; mean CT_MAX_ temperature recorded from species’ independent assays (gold points), and in mixed species control blocks (pink points). There was no global significant difference between CT_MAX_ estimates across all species (Block: F_1,831_ = 3.246, *P* = 0.072). However a significant experimental block*species interaction was found for 4 of the 41 species tested (interaction term: F_40,831_ = 3.7589, P < 0,001); *Drosophila sechelia, D. nasuta, D. yakuba* & *D. tiesseri* all scored significantly higher CT_MAX_ in our mixed species control block than in their own individual CT_MAX_ assays. Right panel: The correlation between estimated CT_MAX_ values from both blocks was strongly positive (coefficient = 1.07, F_1,37_ = 309.84, P < 0.001), and explained 89% of the variation.

**Extended Data Table 1: Correlations between TFL80, LT80, and CT**_**MAX**_. Multiple methods for phylogenetically controlled correlations were used to test the extent to which these three physiological limits are proxies for one another. In all four methods TFL80 correlates less strongly with CT_max_ and LT80 than the two measures of critical limits correlate with each other. Phylogenetic least squares (pgls) allows for estimation of phylogenetic signal (Pagel’s λ) in the residuals of the y-variable, thus correlation coefficients from this method are sensitive to which variable is assigned as the predictor and which as the response. To account for this, we present two pgls() outputs for each pair of traits. Phylogenetic independent contrasts (PIC) essentially assumes a phylogenetic signal of λ = 1 (complete Brownian motion evolution of the trait). Linear models (‘lm’) make no adjustment for non-independence in trait values between closely related species. ‘corphylo’ from the ‘ape’ R package allows for the three-way correlation matrix to be estimated simultaneously and produces estimates of phylogenetic signal (d) of each trait under an Ornstein-Uhlenbeck process. TFL80 here measured 7-days post heat-stress. “-”denotes model estimates are identical to the reciprocal x∼y configuration.

**Extended Data Table 2:** Summaries of phylogenetically controlled models that predict maximum environmental temperature (T_max_) by either CT_MAX_, LT80 or TFL80 independently of each other. Annual precipitation (P_ANN_) was included as an interaction term with physiological limits. adjR^2^ and phylogenetic signal in model residuals (Pagel’s λ) given for final best-fit model derived from AICc model selection and inspection of model residuals. Terms retained after model selection shown in italics. All continuous predictors centred and scaled to the mean.

**Extended Data Table 3:** Summary model fits of species’ safety margins (SM) to absolute latitude (Lat^2^), the physiological limit used in their calculation (‘Limit’), and the interaction between ‘Lat^2^’ and ‘Limit’. ‘Species’ identity is included as a random intercept term. We show models of both “Central safety margin” and the “Distribution Safety Margin” as per Kellermann^8^ Central safety margins are calculated as the difference between the physiological limit and the mean maximum temperature experienced in every know location in the species’ distribution. “Distribution safety margins” capture the environmental conditions at the upper thermal edge of specie’s known distribution by adding 1 standard deviation to the mean maximum temperature. R^2^ given for final best-fit model derived from AICc model selection and inspection of model residuals. Significance of main terms given by Type II sum of squares F-tests with Kenward-Rogers degrees of freedom. Terms retained after model selection shown in italics. Post-hoc Tukey tests were run with the ‘emmeans’ package to identify significant differences between levels of “Limits”. TFL80 measured 7-days post heat shock.

**Extended Data Table 4: Using 50% thresholds to predict distributions**. Summaries of phylogenetically controlled models that predict maximum environmental temperature (T_max_) by either CT_MAX_, LT50 or TFL50 independently of each other. Annual precipitation (P_ANN_) was included as an interaction term with physiological limits. R^2^ and phylogenetic signal in model residuals (Pagel’s λ) given for final best-fit model derived from AICc model selection and inspection of model residuals. Terms retained after model selection shown in italics. All continuous predictors centred and scaled to the mean.

