## Supplementary Methods, Results and Tabls of Species details for "Temperatures that sterilise males better predict global species distributions than lethal temperatures"

**Supplementary Information**

Species Verification:

Where species identity could not be verified through evident morphological traits, we used COI sequence identity to determine phylogenetic position.

We extracted DNA from 2-3 adult male flies with DNeasy kits (Qiagen) following the manufacturer’s invertebrate protocol. We PCR amplified a portion of the mitochondrial universal barcode gene cytochrome oxidase subunit 1 using the primers C1-J-1718 (5’ – GGAGGATTTGGAAATTGATTAGT – 3’) and C1-N-2191 (5’ – CCCGGTAAAATTAAAATATAAACTTC – 3’) using HotStart Taq (Promega) with (5-minute initial heating, 30 cycles at 95°C for 30s, 56 for 30s, and 72°C for 30, with an final elongation step of 72°C for 120s). PCR products were visualised by SYBRSafe-stained gel electrophoresis and cleaned up using Exonuclease I and Shrimp Alkaline Phosphatase incubation per supplier protocol (BioLine). We then used BigDye based sequence reactions with both forward and reverse primers, followed by NaOH & ethanol precipitation clean-up and precipitation before sequences were analyzed with an ABi 3500XL Genetic Analyzer. Forward and reverse sequences for each specie were aligned to derive a consensus sequence. For species with a putative ID based on stock centre or collaborator expertise we assessed the clustering of these sequences with publicly available CO1 sequences from the same species available on the BOLD database (primarily curated genbank sequences). Clustering was performed with the MAFFT server tool, and clustering via a neighbour-joining tree was visualized, all based on default parameters. For species that were unknown (two *repleta*-like species collected from the field in Madeira and Morocco), we performed the same process as above, but included multiple sequences from multiple species in the *repleta* species group – initial BLAST of these sequences on the NCBI database had given an approximate idea of the species ID.

**Testing For lab adaptation:**

We tested for evidence of lab adaptation in thermal traits in the 36 species for which a date of collection was known.

To test if laboratory conditions have selected for low or high thermal limits, we correlated our measurements of TFL80, LT80 and CT_max_ with time in culture using phylogenetically corrected least squares regression (`pgls()` in `caper`). We found no significant association between time in culture and any upper thermal limit (TFL80: slope = 0.003, *P* = 0.801, LT80: slope = 0.003, *P* = 0.629, CT_max_: slope = -0.002, *P* = 0.877, N = 36). To test if laboratory culture causes stabilising selection (i.e. older cultures have more similar traits), we used Breusch-Pagan tests in the package `lmtest` to test for heteroskedasticity in residuals of each upper thermal limit over time. We found a small but significant deviation from homoskedasticity in LT80 over time (χ^2^ = 4.302, df = 1, *P* = 0.038), but no significant deviation in TFL80 (χ^2^ = 0.763, df = 1, *P* = 0.382) nor CTmax (χ^2^ = 2.036, df = 1, *P* = 0.154).

To test if our qualitative result – that TFLs better correlate with species’ thermal environments than lethal temperatures – is a product of lab-adaptation we subset our data into only the ‘newest’ 11 species (all field-collected since 2010) and modelled this subset in the same way described for the main data. In this subset TFL80 measured 7-days post heat stress gave the strongest fits to Tmax compared to either LT80 (105% improvement in adjusted R^2^) or CT_MAX_ (21% improvement in adjusted R^2^).

Whilst these results do not discount laboratory adaptation altogether, they strongly suggest that: 1) between-species variation in upper thermal traits is maintained to a large degree under laboratory conditions; 2) The ecological patterns we see are not a by-product of lab adaptation and are reflected more recently collected species.

**Laboratory species Details:**

| **Supplementary Table 1: Details for species used in this study. All species either *D* = *Drosophila* or *Z* = *Zaprionus*** | | | | | | |
| --- | --- | --- | --- | --- | --- | --- |
| **Species** | **Rearing temp (^o^C)** | **Strain details**  (year collected) | **Multiple**  **lines** | **Food** | **Age at heat (days)** | **Temperature treatments (^o^C)** |
| *D. affinis* | 18 | DSSC Stock #:  14012-0141.14  (2007) | N | M | 7 | 23, 29, 30, 31, 32, 33, 34, 35, 36 |
| *D. albomicans* | 23 | DSSC Stock #: 15112-1751.03  Unknown Age | N | A | 7 | 23, 31, 32, 33, 34, 35, 36 |
| *D. aldrichi* | 23 | Wild caught single line, Texas (2019) | N | A | 7 | 23, 33, 34, 35, 36, 37, 38, 39, 40 |
| *D. americana* | 23 | DSSC Stock #: 15010‑0951.20  (2004) ISOFEMALE | N | M | 7 | 23, 33, 34, 35, 36, 37, 38 |
| *D. ananassae* | 25 | DSSC Stock #: 14024-0371.13  (1945) | N | A | 7 | 25, 30, 33, 34, 35, 36, 37, 38 |
| *D. arizonae* | 23 | DSSC Stock #: 15081-1271-26  (2006) ISOFEMALE | N | B | 7 | 23, 35, 36, 37, 38, 39, 40, 41 |
| *D. biarmipes* | 23 | DSSC Stock #: 14023-0361.11  (2011) | N | A | 7 | 23, 30, 31, 32, 33, 34, 35 |
| *D. bifasciata* | 18 | DSSC Stock #:  14012-0181.02  (2003) ISOFEMALE | N | A | 7 | 18, 25, 27, 29, 30, 31, 32, 33, 34 |
| *D. bipectinata* | 23 | DSSC Stock #:  14024-0381.20  (2011) | N | A | 7 | 23, 30, 31, 32, 33, 34, 35, 36 |
| *D. borealis* | 23 | DSSC Stock #:  15010-0961.10  (2005) ISOFEMALE | N | A | 14 | 23, 31, 32, 33, 34, 35, 36, 37, 38, 39 |
| *D. buzzatii* | 23 | DSSC Stock #:  15081‑1291.02  (1958) | N | M | 7 | 23, 32, 34, 35, 36, 37, 38, 39 |
| *D. eohydei* | 23 | DSSC Stock #:  15085-1631.00  (1956) | N | B | 14 | 23, 31, 32, 33, 34, 35, 36, 37, 38 |
| *D. erecta* | 23 | Cambridge Fly Facility: S-18  (1989) | N | A | 7 | 23, 29, 30, 31, 32, 33, 34, 35, 36, 37, 38, 39 |
| *D. flavomontana* | 23 | DSSC Stock #: 15010-0981  (1949) ISOFEMALE | N | M | 14 | 23, 30, 31, 32, 33, 34, 35, 36, 37 |
| *D. hydei* | 25 | Mixed, long-term lab population at University of Liverpool | Y | B | 14 | 25, 29, 30, 31, 32, 33, 34, 35, 36, 37 |
| *D. lacicola* | 23 | DSSC Stock #: 15010-0991.14  (1949) ISOFEMALE | N | M | 7 | 23, 31, 32, 33, 34, 35, 36, 37, 38 |
| *D. littoralis* | 23 | DSSC Stock #:  15010-1001.03  (1967) | N | B | 14 | 23, 25, 30, 33, 34, 35, 36, 37, 38, 33, 34, 35, 36, 37 |
| *D. lummei* | 23 | DSSC Stock #:  15010-1011.07  (Unknow date) | N | M | 7 | 23, 25, 26, 27, 28, 29, 30, 31, 32, 33, 34, 35, 36, 37 |
| *D. madeirensis* | 18 | Mix of 3 lines, wild caught  (2018) | Y | A | 7 | 18, 27, 28, 29, 30, 31, 32, 33, 34 |
| *D. mayaguana* | 23 | DSSC Stock #:  15081-1397.00  (1985) | N | M | 7 | 23, 35, 36, 37, 38, 39, 40 |
| *D. melanogaster* | 25 | ‘Dahommey’ outbred lab population | Y | A | 7 | 25, 30, 31, 32, 33, 34, 35, 36, 37, 38 |
| *D. mercatorum* | 23 | Wild-caught Madeira  (2018) single line | N | A | 7 | 23, 31, 32, 33, 34, 35, 36, 37, 38, 39, 40 |
| *D. micromelanica* | 23 | DSSC Stock #:  15030-1151.01  (unknown date) | N | A | 7 | 23, 30, 31, 32, 33, 34, 35, 36, 37, 38 |
| *D. melanica* | 23 | DSSC Stock #:  15030-1141.03  (unknown date) | N | A | 7 | 23, 32, 33, 34, 35, 36, 37, 38 |
| *D. mojavensis* | 23 | DSSC Stock #: 15081-1352.00  (unknown date) | Y | B | 7 | 23, 34, 35, 36, 37, 38, 39, 40, 41 |
| *D. montana* | 18 | Mixed population of IsoLines in ^41^  (2013) | Y | M* | 21 | 18, 31, 32, 33, 34, 35, 36, 37 |
| *D. nasuta* | 23 | DSSC Stock #:  15112-1781.13  (2004) ISOFEMALE | N | A | 7 | 233, 30, 31, 32, 33, 34, 35, 36, 37 |
| *D. novamexicana* | 23 | DSSC Stock #: 15010-1031.04  (1949) ISOFEMALE | Y | B | 7 | 23, 28, 30, 32, 33, 34, 35, 36, 37, 38 |
| *D. obscura* | 23 | Mixed population collected in UK (2012) | Y | P | 7 | 23, 28, 29, 30, 31, 32, 33 |
| *D. paranaensis* | 23 | DSSC Stock #:  15082-1541.09  (2004) | N | A | 7 | 23, 31, 32, 33, 34, 35, 35, 37 |
| *D. pseudoobscura* | 18 | Mixed population IsoLines in ^42^  (2008) | Y | A* | 7 | 23, 29, 30, 31, 32, 33, 34, 35 |
| *D. repleta* | 23 | DSSC Stock #:  15084-1611.01  (1953) | N | A | 7 | 23, 30, 31, 32, 33, 34, 35, 36, 38 |
| *D. santomea* | 23 | DSSC Stock #: 14021-0271  (1998) ISOFEMALE | N | A | 7 | 23, 29, 30, 31, 32, 33, 34, 35, 36, 37 |
| *D. sechellia* | 23 | Cambridge Fly Facility: S-32  (1995) | Y | P | 7 | 23, 29, 30, 31, 32, 33, 34, 35, 36, 37 |
| *D. simulans* | 25 | Mixed population of lines collected across Eastern Europe (2016) | Y | A | 7 | 25, 30, 32, 33, 34, 35, 36, 37 |
| *D. subobscura* | 18 | Mixed population from ISOFEMALE lines in ^43^  (2013 & 2016) | Y | A | 7 | 18, 25, 27, 29, 30, 31, 32, 33, 34 |
| *D. suzukii* | 23 | Mixed populations of 3 IsoLines caught in Madeira (March 2018) and single lines each from the USA and Japan | Y | A | 7 | 23, 27, 29, 30, 31, 32, 33, 34, 35, 36, 37 |
| *D. takahashii* | 23 | DSSC Stock #: 14022-0311.10  (2005) | N | A | 7 | 23, 30, 31, 32, 33, 34, 35, 36, 37 |
| *D. teissieri* | 23 | DSSC Stock #:  14021‑0257.01  (unknown date) | N | A | 7 | 23, 28, 29, 30, 31, 32, 33, 34, 35, 36, 37 |
| *D. virilis* | 23 | Cambridge Fly Facility: S-4  (1991) | Y | P | 7 | 23, 26, 28, 30, 32, 34, 35, 36, 37, 38 |
| *D. yakuba* | 23 | Cambridge Fly Facility: S-15  (1973) | Y | A | 7 | 23, 30, 31, 32, 33, 34, 35, 36, 37, 38 |
| *Zaprionus indianus* | 23 | DSSC Stock #: 50001-0001.05  (2004) ISOFEMALE | N | B | 7 | 23, 33, 34, 35, 36, 37, 38, 39 |
| *Zaprionus tuberculatus* | 23 | Wild caught  (2019) | Y | B | 7 | 23, 27, 29, 31, 32, 33, 34, 35, 36, 37, 38 |
