## Extended Data Figure 1 - 6, Extended Data Tables 1 - 4 for "Temperatures that sterilise males better predict global species distributions than lethal temperatures"

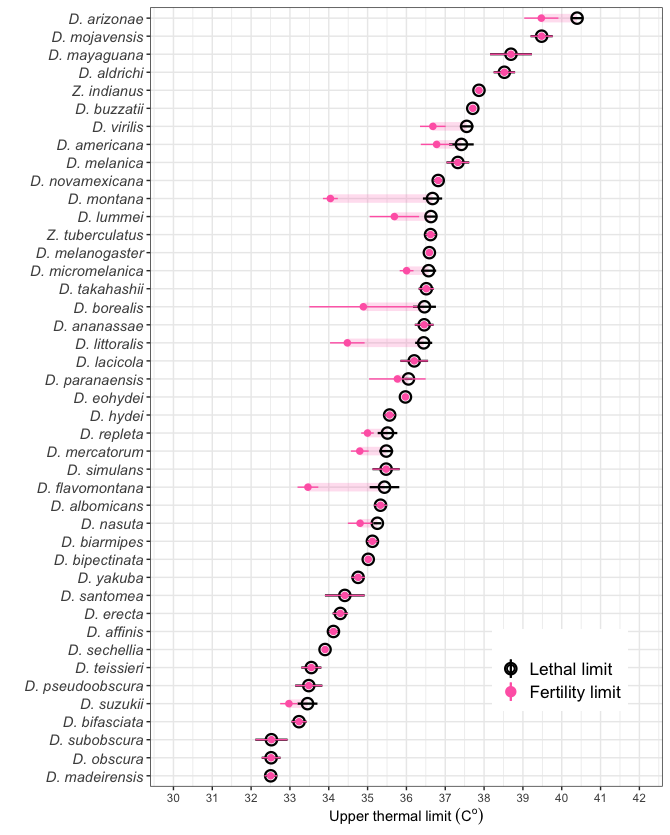

**Extended Data Figure 1: TFL80 and LT80 of Drosophila immediately after heat stress.** Several species of Drosophila lose 80% fertility at cooler-than-lethal temperatures immediately following heat-shock. LT80 (black circles) and TFL80 (pink circles). Errors for both measures are 95% confidence intervals generated from dose response model estimates. Fertility loss measured as ability to sire any offspring between 1 – 6 days post heat-stress.

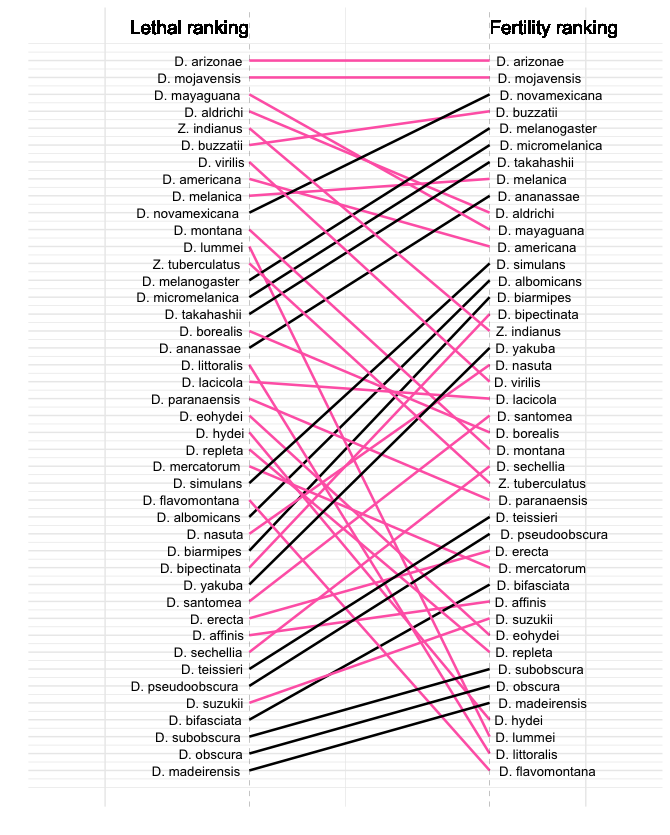

**Extended Data Figure 2: Relative ranking of species thermal robustness based on LT80 and TFL80.** Most to least heat tolerant species ranked from top to bottom. Pink bars indicate species with significantly lower TFL80 than LT80. Data based on TFL80 recorded 7-days post-heat shock.

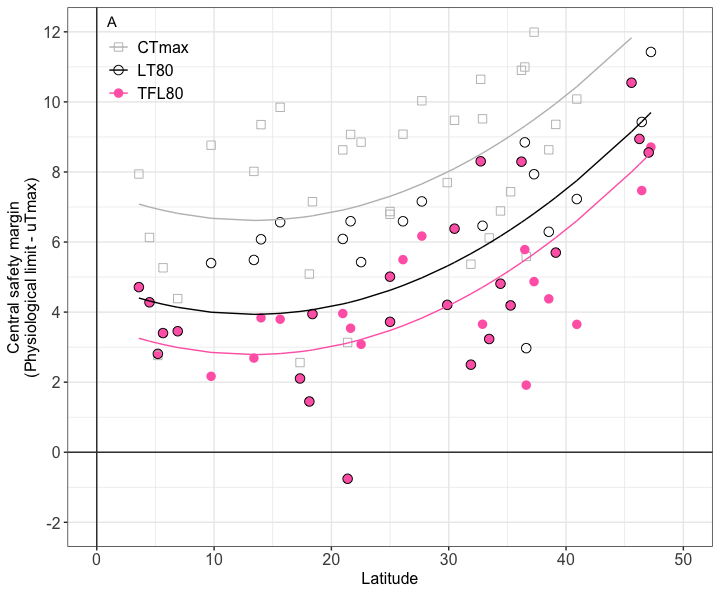

**Extended Data Figure 3: Central Thermal safety margins for 43 Drosophila species.** Margins calculated using the physiological limit minus the mean maximum temperature across all recorded locations for each species. Grey squares and line are safety margins calculated using CT_MAX_ values, black circles and lines are based on LT80, and pink points and lines are based on TFL80 measured 7-days after heat stress. The three physiological limits produce significantly different safety margin estimates (Central safety margins: lmer: F_2_ = 28.23, *P* < 0.001, estimate = -1.17). TFL80 measured at 7 days post-heat stress predicts mean central safety margins of 4.97 ± 2.42^o^C. This is significantly smaller than margins based on CT_MAX_ (mean central safety margins: 8.81 ± 3.26^o^C, pairwise Tukey tests P<0.001) and LT80 (central safety margins: 6.13 ± 3.09^o^C, pairwise Tukey tests P<0.001). Latitude shown here is the mean absolute latitude i.e. the sign of negative latitudes has been removed to give relative distance from the equator. Fitted lines are predictions from models in which the limit type (TFL, LT or CT_MAX_) and latitude^2^ are fixed effects and species identity is a random effect to account for repeated measures across species (see Extended Data Table 3).

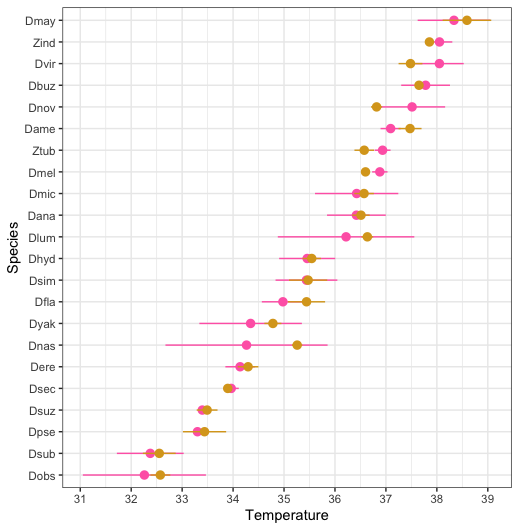

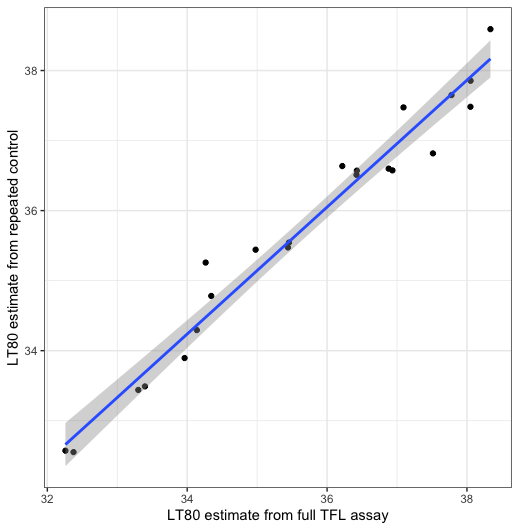

**Extended Data Figure 4: Repeatability of LT80**. Repeatability was high across the 22 *Drosophila* species tested. Left panel: There was no significant difference in the estimate of LT80 between full runs and control repeats in any of these species (significance measured as non-overlapping CIs of point estimates of LT80). Pink points = estimate from single species assay, gold point = estimate from simultaneous multispecies control assay. Right panel: The correlation between the two independent measurements of LT80 across these species was strongly positive and explained a high degree of variation in the data (coefficient = 1.06, R^2^ = 0.96).

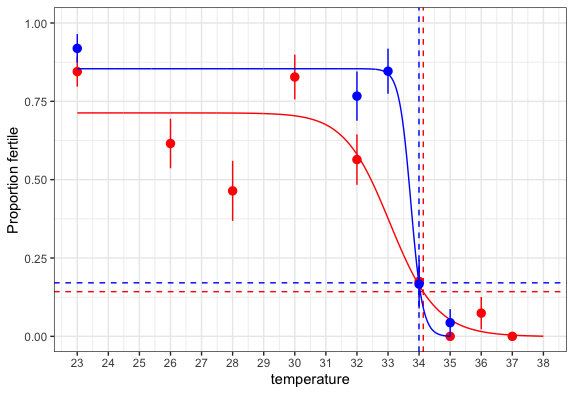

**Extended Data Figure 5: Repeatability of TFL80 for *Drosophila virilis*.** We independently repeated the TFL80 assay for one species in which we found a significant difference between LT80 and TFL80. Two independent runs of sexually mature males were conducted by two researchers 6 months apart. Red = original data used to calculate TFL80 for *D. virilis* in this manuscript, blue = repeated assay. Dashed lines intersect at 80% threshold. Line fits predicted by 3-parameter dose-response model. The fly stock and equipment were identical.

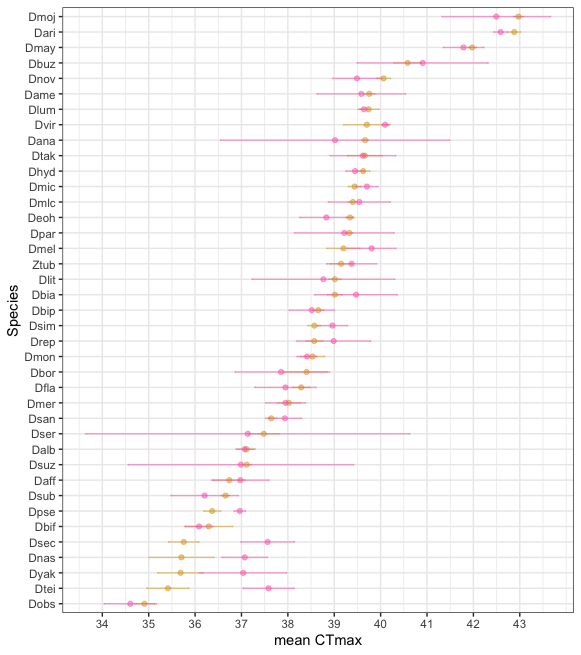

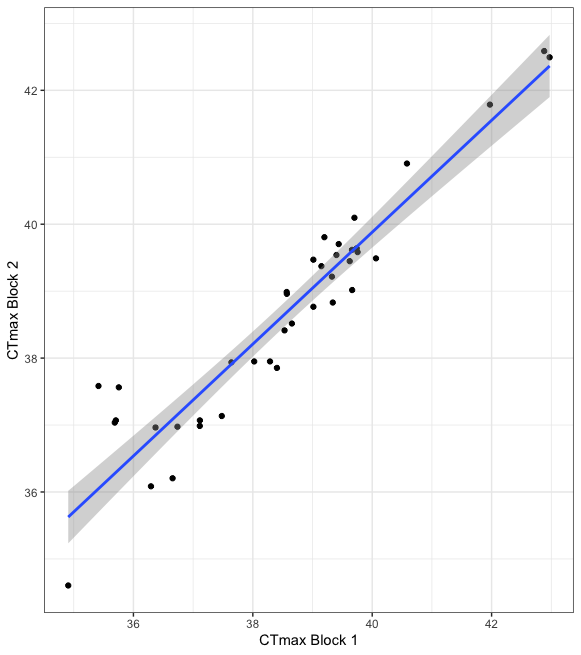

**Extended Data Figure 6: Repeatability of knockdown CT_MAX_.** Left panel; mean CT_MAX_ temperature recorded from species’ independent assays (gold points), and in mixed species control blocks (pink points). There was no global significant difference between CT_MAX_ estimates across all species (Block: F_1,831_ = 3.246, *P* = 0.072). However a significant experimental block*species interaction was found for 4 of the 41 species tested (interaction term: F_40,831_ = 3.7589, P < 0,001); *Drosophila sechelia, D. nasuta, D. yakuba* & *D. tiesseri* all scored significantly higher CT_MAX_ in our mixed species control block than in their own individual CT_MAX_ assays. Right panel: The correlation between estimated CT_MAX_ values from both blocks was strongly positive (coefficient = 1.07, F_1,37_ = 309.84, P < 0.001), and explained 89% of the variation.

| **Extended Data Table 1:** Correlations between TFL80, LT80, and CT_MAX_. Multiple methods for phylogenetically controlled correlations were used to test the extent to which these three physiological limits are proxies for one another. In all four methods TFL80 correlates less strongly with CT_MAX_ and LT80 than the two measures of critical limits correlate with each other. Phylogenetic least squares (pgls) allows for estimation of phylogenetic signal (Pagel’s λ) in the residuals of the y-variable, thus correlation coefficients from this method are sensitive to which variable is assigned as the predictor and which as the response. To account for this, we present two pgls() outputs for each pair of traits. Phylogenetic independent contrasts (PIC) essentially assumes a phylogenetic signal of λ = 1 (complete Brownian motion evolution of the trait). Linear models (‘lm’) make no adjustment for non-independence in trait values between closely related species. `corphylo` from the `ape` R package allows for the three-way correlation matrix to be estimated simultaneously and produces estimates of phylogenetic signal (d) of each trait under an Ornstein-Uhlenbeck process. TFL80 here measured 7-days post heat-stress. “-“denotes model estimates are identical to the reciprocal x~y configuration. | | | | | | | | | |
| --- | --- | --- | --- | --- | --- | --- | --- | --- | --- |
|  | PGLS  (ML λ) | | | PICs  (λ = 1) | | lm  (no accounting for phylogeny) | | Corphylo | |
| Model (y ~ x) | Corr | Sig | Phylo signal (λ) | Corr | Sig | Corr | Sig | Corr | Phylo signal (d) |
| CT_MAX_ ~ LT80 | 0.850 | *** | 0.341 | 0.560 | *** | 0.911 | *** | 0.702 | CTmax = 0.450,  LT80 = 0.592,  TFL80 = 0.238 |
| LT80 ~ CT_MAX_ | 0.820 | *** | 0.570 | - | - | - | - | - | - |
| TFL80 ~ LT80 | 0.734 | *** | 0.570 | 0.546 | *** | 0.691 | *** | 0.654 | - |
| LT80 ~ TFL80 | 0.705 | *** | 0.833 | - | - | - | - | - | - |
| TFL80 ~ CT_MAX_ | 0.567 | *** | 0.543 | 0.397 | ** | 0.590 | *** | 0.509 | - |
| CT_MAX_ ~ TFL80 | 0.534 | *** | 0.762 | - | - | - | - | - | - |

| **Extended Data Table 2:** Summaries of phylogenetically controlled models that predict maximum environmental temperature (T_max_) by either CT_MAX_, LT80 or TFL80 independently of each other. Annual precipitation (P_ANN_) was included as an interaction term with physiological limits. adjR^2^ and phylogenetic signal in model residuals (Pagel’s λ) given for final best-fit model derived from AICc model selection and inspection of model residuals. Terms retained after model selection shown in italics. All continuous predictors centred and scaled to the mean. | | | | | | | |
| --- | --- | --- | --- | --- | --- | --- | --- |
| **Full Model** | **Terms** | **Estimate**  **(se)** | **DF** | **t** | **P** | **^(adj)^R^2^** | **Pagel’s λ** |
| T_max_~ CT_max_ + (CT_max_:P_ANN_) | *CT_max_ : P_ANN_* | *1.500*  *(0.566)* | *40* | *2.647* | *0.012* | 0.186 | 0.375 |
|  | *CT_max_ : P_ANN_* | *-0.925*  *(0.445)* | *40* | *-2.077* | *0.044* |  |  |
| T_max_~ LT80 + (LT80:P_ANN_) | *LT80* | *2.073*  *(0.617)* | *41* | *3.360* | *0.002* | 0.197 | 0.441 |
|  | LT80 : P_ANN_ | -0.553  (0.544) | 40 | -1.018 | 0.315 |  |  |
| T_max_~ TFL80 + (TFL80:P_ANN_) (Immediately post-heat) | *TFL80* | *2.218*  *(0.525)* | *41* | *4.225* | *<0.001* | 0.286 | 0.361 |
|  | TFL80 : P_ANN_ | -0.181  (0.500) | 40 | 0.361 | 0.719 |  |  |
| T_max_~ TFL80 + (TFL80:P_ANN_)  (7 days post-heat) | *TFL80* | *2.081*  *(0.415)* | *41* | *5.014* | *<0.001* | 0.365 | 0.281 |
|  | TFL80 : P_ANN_ | -0.342  (0.470) | 40 | -0.730 | 0.469 |  |  |

| **Extended Data Table 3:** Summary model fits of species’ safety margins (SM) to absolute latitude (Lat^2^), the physiological limit used in their calculation (‘Limit’), and the interaction between ‘Lat^2^‘ and ‘Limit’. ‘Species’ identity is included as a random intercept term. We show models of both “Central safety margin“ and the “Distribution Safety Margin” *as per* Kellermann^8^. Central safety margins are calculated as the difference between the physiological limit and the mean maximum temperature experienced in every know location in the species’ distribution. “Distribution safety margins” capture the environmental conditions at the upper thermal edge of specie’s known distribution by adding 1 standard deviation to the mean maximum temperature. R^2^ given for final best-fit model derived from AICc model selection and inspection of model residuals. Significance of main terms given by Type II sum of squares F-tests with Kenward-Rogers degrees of freedom. Terms retained after model selection shown in italics. Post-hoc Tukey tests were run with the `emmeans` package to identify significant differences between levels of “Limits”. TFL80 measured 7-days post heat shock. | | | | | | | |
| --- | --- | --- | --- | --- | --- | --- | --- |
| Response Variable | Fixed Effects | DF | F | P | R^2^ | Random Effect  (Species ID) | Tukey contrasts *P* |
| Central Safety Margin  (Limit - μT_max_) | Lat^2^*Limit | 2,80 | 0.710 | 0.494 | 0.70 | 2.513 | All <0.001 |
|  | *Lat^2^* | *1,40* | *163.92* | *<0.001* |  |  |  |
|  | *Limit* | *2,84* | *16.56* | *<0.001* |  |  |  |
| Distribution Safety Margin  (Limit – [μT_max_ + 1SD]) | Lat^2^*Limit | 2,80 | 0.710 | 0.494 | 0.57 | 5.050 | All <0.001 |
|  | *Lat^2^* | *1,40* | *163.92* | *0.001* |  |  |  |
|  | *Limit* | *2,84* | *12.19* | *<0.001* |  |  |  |

| **Extended Data Table 4: Using 50% thresholds to predict distributions.** Summaries of phylogenetically controlled models that predict maximum environmental temperature (T_max_) by either CT_MAX_, LT50 or TFL50 independently of each other. Annual precipitation (P_ANN_) was included as an interaction term with physiological limits. R^2^ and phylogenetic signal in model residuals (Pagel’s λ) given for final best-fit model derived from AICc model selection and inspection of model residuals. Terms retained after model selection shown in italics. All continuous predictors centred and scaled to the mean. | | | | | | | |
| --- | --- | --- | --- | --- | --- | --- | --- |
| **Full Model** | **Terms** | **Estimate**  **(se)** | **DF** | **t** | **P** | **^(adj)^R^2^** | **Pagel’s λ** |
| T_max_~ CT_max_ + (CT_max_:P_ANN_) | *CT_max_ : P_ANN_* | *1.500*  *(0.566)* | *40* | *2.647* | *0.012* | 0.186 | 0.375 |
|  | *CT_max_ : P_ANN_* | *-0.925*  *(0.445)* | *40* | *-2.077* | *0.044* |  |  |
| T_max_~ LT50 + (LT50:P_ANN_) | *LT50* | *2.050*  *(0.605)* | *41* | *3.388* | *0.002* | 0.200 | 0.435 |
|  | LT50 : P_ANN_ | -0.718  (0.524) | 40 | -1.371 | 0.178 |  |  |
| T_max_~ TFL50 + (TFL50:P_ANN_) (Immediately post-heat) | *TFL50* | *2.468*  *(0.468)* | *41* | *5.273* | *<0.001* | 0.333 | 0.390 |
|  | TFL50 : P_ANN_ | -0.556  (0.480) | 40 | -1.161 | 0.252 |  |  |
| T_max_~ TFL50 + (TFL50:P_ANN_)  (7 days post-heat) | *TFL50* | *1.944*  *(0.506)* | *40* | *3.841* | *<0.001* | 0.470 | 0.000 |
|  | *TFL50 : P_ANN_* | *-1.210*  *(0.453)* | *40* | *-2.672* | *0.011* |  |  |
